# Contamination of widely used databases compromises rRNA based taxonomic assignment of metagenomes and metatranscriptomes

**DOI:** 10.64898/2026.07.22.739903

**Authors:** Alastair Grant, Charli S Davies

## Abstract

Taxonomic annotation of metagenomic and metatranscriptomic datasets frequently relies on aligning small subunit (SSU) rRNA reads to annotated reference databases. However, widely used SSU databases, including SILVA, GTDB, Eukaryome, and those distributed with SortMeRNA, are contaminated with large subunit (LSU) rRNA sequences. Low levels of LSU contamination can lead to large numbers of LSU reads being incorrectly identified as SSU and have serious consequences for downstream taxonomic assignments. The problem can be avoided by rigorously removing LSU sequences from databases used for taxonomic annotation, which we have done for KSGP 4.0. Alternatively, initial SSU read selection can be carried out with a carefully curated small SSU database that is free of LSU contamination. We illustrate these two approaches in combination using a metatranscriptomic dataset from an estuarine sediment. Without database cleaning, LSU-derived reads can make up half of supposed SSU sequences and are assigned to a small number of apparently dominant but artefactual taxa. Database cleaning removes this problem and we provide a script, total_rnaseq, to annotate total RNASeq or metagenomic data using this approach. The KSGP 4.0 database provides substantially improved annotation of Archaea compared to the SILVA database and moderate and small improvements for eukaryotes and bacteria respectively.

## Introduction

Metagenomics and metatranscriptomics provide information on the abundances of all organisms present in a sample without the taxonomic biases intrinsic to PCR based approaches (1-6). In principle, all retrieved sequences are taxonomically informative and can be annotated using databases of whole genomes (7,8). But full genome sequences are available for only a small proportion of organisms, a problem that is particularly acute for eukaryotic microbes (9). Therefore it is common to base taxonomic annotations of metagenomes on small subunit ribosomal RNA sequences (SSU) for which very large databases of taxonomically characterised sequences are available (10,11). This approach is also central to metatranscriptomics of total RNA (without rRNA removal), also called Total RNASeq, double RNASeq or community RNASeq (1,12-20). In both cases, individual DNA or RNA reads are compared to a database of taxonomically annotated sequences. Taxonomic assignment may use Bayesian methods as in RDP (21); kmer matching as in Sintax (22), hidden Markov models (23) or Lowest Common Ancestor (LCA) assignments based on sequence alignments or kmer matches as in Kraken and LotuS2 (8,24). Database searching may be carried out for all reads passing a quality threshold, or putative SSU reads may first be extracted using an algorithm that is faster than full database searching, such as SortMeRNA (25) or barrnap (26).

SSU databases such as SILVA (27,28), RDP (21), PR2 (29,30), Eukaryome (31) and KSGP (32) have been developed primarily for the analysis of metabarcoding data, where all query sequences are derived from the target region. However, metagenomic and metatranscriptomic data contain sequences from other kinds of RNA. In particular, most SSU and LSU rRNA genes are organised into operons and transcribed together (33-35) so we would expect approximately equimolar amounts of SSU and LSU molecules in environmentally derived DNA and RNA. But as the LSU molecule is larger, LSU reads are likely to be more numerous than SSU reads in metagenomic and metatranscriptomic data.

Commonly used SSU databases are known to include numbers of incorrectly annotated sequences (32,36,37). Contamination of SSU databases with LSU sequences could have serious consequences for taxonomic assignment of metagenomes and metatranscriptomes, as LSU reads in the data may be wrongly identified as SSU. The potential for this has been noted previously and the phyloFlash pipeline uses a cleaned version of the SILVA database in response (38). But the full significance of this problem is not widely recognised and use of databases without LSU removal remains common.

Here we show that LSU contamination of SSU databases in widespread use can lead to LSU reads contributing up to half of those identified as SSU, seriously impacting the integrity of taxonomic assignments. We identify strategies to circumvent this challenge and provide a script that processes raw metagenomic or total metatranscriptomic reads and generates tables of read counts at all taxonomic levels.

## Methods

### Sample collection, RNA extraction and sequencing

Intertidal mud was collected from Restronguet Creek, Cornwall, SW England (50.20777° N 5.10621° W, site RA in reference 39). This is a low salinity site (6.6 when sampled) that is severely contaminated with Cu, Zn, As and other metals. DNA and RNA were extracted using the RNeasy PowerSoil Total RNA kit and DNA elution accessory kit (Qiagen, Hilden, Germany) according to an optimised version of the manufacturer’s instructions with an added heat block step (45°C for 15 minutes) prior to the solution being added to the column. Nucleic acids were quantified using Invitrogen Qubit RNA and dsDNA broad range kits (ThermoFisher, Loughborough, UK) measuring fluorescence with either a qPCR machine or a Qubit 4 fluorimeter. After removing DNA contamination using the Invitrogen (Waltham, MA) DNA-free kit, following manufacturer’s instructions, 150PE total RNA sequencing, without ribosomal RNA depletion, was carried out on an Illumina NovaSeq X plus instrument by a commercial service provider (Novogene, Cambridge, UK).

### Processing RNASeq data

Sequencing adapters and low quality bases were removed using Trimmomatic (47). Overlapping read pairs were merged using USEARCH fastq mergepairs (48). Where reads were not successfully merged only the forward (R1) was retained because of concerns about the impact of concatenation of unmerged reads on the percentage similarity of local alignments used in our approach to taxonomic assignment. These were combined with surviving reads from pairs where only one read survived quality trimming.

### Databases used

Databases used or constructed during this study are detailed in Table 1. Manually curated SSU and LSU sequences for prokaryotes and eukaryotes are available from Rfam (43); prokaryote and fungal SSU sequences (but not other eukaryotes) from RefSeq (40,41) and prokaryote LSU and eukaryote SSU and LSU from the NCBI BLAST databases (42). NCBI Prokaryote LSU; Eukaryote LSU and SSU rRNA BLAST databases were chosen in preference to Rfam as they contain representatives of more taxa and can be downloaded directly, but the Rfam database was used for 5.8S sequences.

**Table 1.** Sequence databases used in this study. ^*^Indicates that database is available on the GitHub page associated with this paper (https://github.com/AGrantUEA/total_RNASeq).

| Database | Notes | Reference |
| --- | --- | --- |
| NCBI SSU* | NCBI Eukaryote BLAST database dated 26 April 2024 concatenated with Bacteria and Archaea 16S sequences from the REFSEQ targeted loci project dated 21 February 2025 | (40-42) |
| NCBI LSU* | NCBI prokaryote and eukaryote LSU BLAST databases dated 26 April 2024. Except for the initial comparison with NCBI SSU, a single accession contaminated with SSU rRNA was removed (see results). | (42) |
| Rfam 5.8S | Downloaded 23 September 2025 | (43) |
| SortMeRNA<br>SSU<br>Sensitive | SSU Sensitive database version 4.3 | (25) |
| SortMeRNA<br>cleaned* | SSU Sensitive database after removing accessions containing LSU sequence using SortMeRNA and the NCBI LSU database. | See<br>methods |
| SILVA 138.1<br>SSU Ref<br>NR99 | Using standardised taxonomy distributed with LotuS2 | (24,27,44) |
| SILVA<br>cleaned* | SILVA 138.1 after removing all sequences identified as containing LSU sequence using USEARCH with the NCBI LSU database | See<br>methods |
| GTDB r226<br>SSU | 16S sequences from GTDB r226 | (45) |
| Eukaryome<br>2.0 SSU | SSU subset of Eukaryome 2.0 as distributed by authors – sequences are not truncated at 3' end of 18S | (31) |
| Eukaryome<br>2.0 SSU<br>trimmed* | 18S sequences from Eukaryome SSU after removal of prokaryotes, mitochondria and plastids and truncated at 3' end using ITSx. | See<br>methods |
| Midori2<br>version 267 | Contributes mitochondrial SSU sequences to KSGP 4.0 | (46) |
| PR2 5.0 | Used as source of plastid SSU sequences in KSGP 4.0 | (29,30) |
| GTDB+ 4.0 | Component part of KSGP. Cleaned prokaryote SSU sequences from GTDB after removing taxonomic inconsistencies plus SSU sequences from Eukaryome, PR2 and Midori as described above. | (32) |
| KSGP 4.0 | GTDB+ combined with long environmental SSU rRNA sequences and prokaryote SSU sequences from SILVA not present in GTDB+. | (32) |

### Construction of version 4.0 of the GTDB+ and KSGP databases

GTDB+, which forms part of our KSGP database, is a cleaned database of SSU sequences from prokaryotes, eukaryotes, mitochondria and plastids. KSGP construction is described elsewhere (32), but in the updated version 4.0, eukaryote 18S sequences have been obtained from the Eukaryome database as this is more comprehensive than the PR2 database used previously and we have also increased the rigour of sequence screening. In brief; 16S sequences from the GTDB database are screened using Ribovore (49), removing those assigned to the incorrect domain or labelled as fail or pass with unexpected features. This is followed by two rounds of RDP LOOT (leave one out, 50), the first retaining only those assigned to the correct domain followed by a second round retaining only those assigned to the correct Class.

A substantial portion of sequences in the Eukaryome SSU database extend beyond the 3’ end of the 18S gene and include part or all of the ITS1, 5.8S, ITS2 and 28S sequences. 18S sequences were extracted from these using ITSx version 1.1.3 (51). Most sequences for which the boundary between the 18S and ITS1 sequences is not detected represent genuine but partial 18S sequences terminated before their 3’ end. These were retained in the database if identified as Eukaryote or Apicomplexan SSU sequences by Ribovore. Sequences containing 5.8S contamination were identified and removed using SortMeRNA and the Rfam 5.8S database.

Plastid sequences were obtained from PR2 and mitochondrial sequences from Midori2, removing any that Ribovore does not assign with confidence as plastids or mitochondria respectively. Plastid sequences in the PR2 and SILVA databases appeared to include a number of chimeric and mis-labelled sequences, so only those chloroplast sequences derived from whole plastid genome sequences were retained. Dinoflagellate chloroplast genomes are much reduced and can contain unusual SSU rRNA sequences (52-54). PR2 5.0 does not include any dinoflagellate SSU sequences derived from whole plastid genomes, so plastid SSU rRNA sequences from *Lepidodinium chlorophorum* (Gymnodiniales; NC_027093) and *Kryptoperidinium foliaceum* (Peridiniales; NC_014267), representing the two largest lineages, were added manually. Finally, any remaining LSU or 5.8S contamination was removed using SortMeRNA as described in the final paragraph of this methods section. KSGP 4.0 (and the GTDB+ subset) are available at https://ksgp.earlham.ac.uk/.

### Assessment of LSU contamination

LSU contamination of SSU databases, including the SILVA; Eukaryome and SortMeRNA databases and/or SSU contamination of LSU databases was initially identified using local searches in USEARCH (55) with a number of pairs of SSU and LSU databases. Likely consequences of this for total RNASeq and rRNA based taxonomic assignment in metagenomics were assessed by splitting sequences in the RefSeq SSU and LSU databases into overlapping 150 bp fragments, starting at 75 bp intervals supplemented by the final 150bp of each sequence where this was not already included. When used with a low similarity threshold and queries that are much shorter than database sequences (as is typical in RNASeq), the default settings of USEARCH (and other rapid alternatives to BLASTN) can terminate a search on a query without finding a match or return a relatively low similarity match, even when a high similarity match is present in the database (A. Grant, pers. Obs.). Heuristics in USEARCH can be disabled by setting maxaccepts and maxrejects to zero. This increases run times very substantially so is not feasible for the analysis of real RNAseq datasets. For the example dataset examined here we used USEARCH with values of 100 for maxaccepts, 5000 for maxrejects and 200 for maxhits. Analysis of a sample of 100 000 reads showed that these parameters found approximately 99.5% of the matches identified by an exhaustive USEARCH (turning off heuristics by setting maxaccepts and maxrejects to zero). This issue will be discussed in more detail elsewhere (A. Grant, in prep).

### Selection of SSU sequences and/or removal of LSU contamination

LSU sequences were removed from SILVA 138.1 using USEARCH and from the SortMeRNA sensitive SSU database and the final versions of GTDB+ and KSGP 4.0 using SortMeRNA, in all cases using a database consisting of concatenated Fasta files extracted from the NCBI Prokaryote and Eukaryote LSU BLAST databases after first removing a single chimeric sequence (see results) and the Rfam 5.8S database. The cleaned databases are referred to below as SILVA-clean and SortMeRNA-clean, while SILVA and SortMeRNA sensitive are used to refer to the original versions. SSU sequences were extracted from sequencing data either with mTAGs using its default hidden Markov models (HMMs) (10) or SortMeRNA using either the NCBI SSU database, the SortMeRNA-cleaned database, or these two databases concatenated (see Table 1).

### Taxonomic assignment

Total and extracted SSU reads were aligned to the KSGP 4.0, SILVA and SILVA-cleaned databases using USEARCH and taxonomy assigned using version 0.25 of the LCA tool from LotuS2 (24). This was carried out separately for all three domains for SSU and LSU reads extracted by mTAGs. Taxonomic assignment based on BLAST searches or equivalent combined with LCA appears to be robust in the face of database heterogeneity (56,57).

## Results and discussion

### Identification of contamination of SSU databases with LSU sequences

Comparison of SSU and LSU databases using tools such as BLASTN or USEARCH identifies cross matches which occur irrespective of which is used as queries and which as the target database. For example, between 90 and 100% of supposed SSU queries had matches in the SortMeRNA LSU database (Table 1). These could result from the presence in one database of sequences that span both SSU and LSU; SSU/LSU chimeras; mis-labelling of SSU sequences as LSU or vice versa. Automated identification of where the contamination lies requires SSU and LSU databases that are entirely free of contamination. This was not true of any of the databases that we examined, but the NCBI SSU and LSU databases showed only a very low level of cross matching (Table 2). Two out of 36 101 SSU sequences have a match in the combined eukaryote and prokaryote LSU database, while one out of 10 622 LSU sequences has a match in the SSU database. Manual examination of these sequences indicates that the cross-matching LSU sequence is chimeric and contains small amounts of SSU sequence. When this was removed from the LSU database, USEARCH no longer detected any matches. The more sensitive SortMeRNA did identify some cross matches, but manual inspection of these showed that the length of the alignment over which matching occurred was short (details not shown).

**Table 2.** Percentages of database sequences with USEARCH matches in the SortMeRNA LSU sensitive database and a combined prokaryote and eukaryote NCBI LSU BLAST database. ^†^Comparison between NCBI SSU and NCBI LSU carried out including SSU contaminant sequence in NCBI LSU database; other comparisons with NCBI LSU exclude this.

| Database used as query | Matches in SortMeRNA LSU sensitive | Matches in NCBI LSU |
| --- | --- | --- |
| NCBI SSU | 100% | 0.006%<sup>†</sup> |
| SortMeRNA SSU Sensitive | 99.9% | 0.06% |
| Silva 138.1 SSU | 99.9% | 0.08% |
| GTDB r226 SSU | 89.8% | 0.014% |
| Eukaryome 2.0 SSU | 98.4% | 54.7% |
| Eukaryome 2.0 SSU trimmed | 99.9% | 0.0003% |

### Quantifying the scale of contamination of SSU databases with LSU sequences

After removing a single chimeric sequence (see above), this small NCBI LSU database can be used to assess the extent of LSU contamination in SSU databases. Even with a low stringency search (70% identity and minimum alignment length of 100 bp) the proportion of sequences in the SILVA, GTDB and SortMeRNA sensitive databases that were contaminated with LSU was very small (0.01% to 0.08%; Table 2) but this rose to 54.7% for the Eukaryome SSU database. The last figure reflects the presence within Eukaryome of long reads spanning a substantial part of the rRNA operon and containing SSU, ITS and LSU sequence. The SSU component of Eukaryome includes all sequences that cover part of SSU without trimming, to avoid the potential for trimmed and non-trimmed versions of the same sequence to be given different taxonomic assignments (Leho Tedersoo, personal communication). When the Eukaryome SSU database was trimmed as described in the methods section, only one sequence (0.0003%) had a match in the NCBI LSU database (Table 2).

When broken into 150 bp fragments, no LSU fragments were matched to the NCBI SSU database and only 5 out of 709 034 SSU fragments were matched to the NCBI LSU database (Table 2), so the LSU database with one sequence removed and the NCBI SSU database were used in subsequent analyses.

### Potential impacts of LSU contamination of SSU databases on rRNA based taxonomic assignments

Although only a small number of SSU sequences in SILVA, GTDB and the SortMeRNA sensitive database are contaminated with LSU sequences, even small amounts of LSU contamination can lead to a large proportion of LSU reads being mis-classified as SSU by both SortMeRNA and USEARCH (Table 3). SortMeRNA classifies between 31.7% and 98% of LSU fragments as SSU, depending upon the database used, with slightly lower numbers for USEARCH. Adding just a single full-length eukaryote or prokaryote LSU molecule to the NCBI SSU database led to misclassification of 35.8% of eukaryote LSU fragments and 33.0% of prokaryote fragments respectively. This does not present a problem when SortMeRNA is being used for its original purpose of removing rRNA sequences so that mRNA sequences can be studied in detail. But small numbers of contaminating LSU sequences in databases could seriously compromise the integrity of taxonomic assignments using SSU sequences. When this occurs, large numbers of LSU fragments may match small numbers of database contaminants and taxa represented by them may dominate the final taxonomic assignments (see below).

**Table 3.**
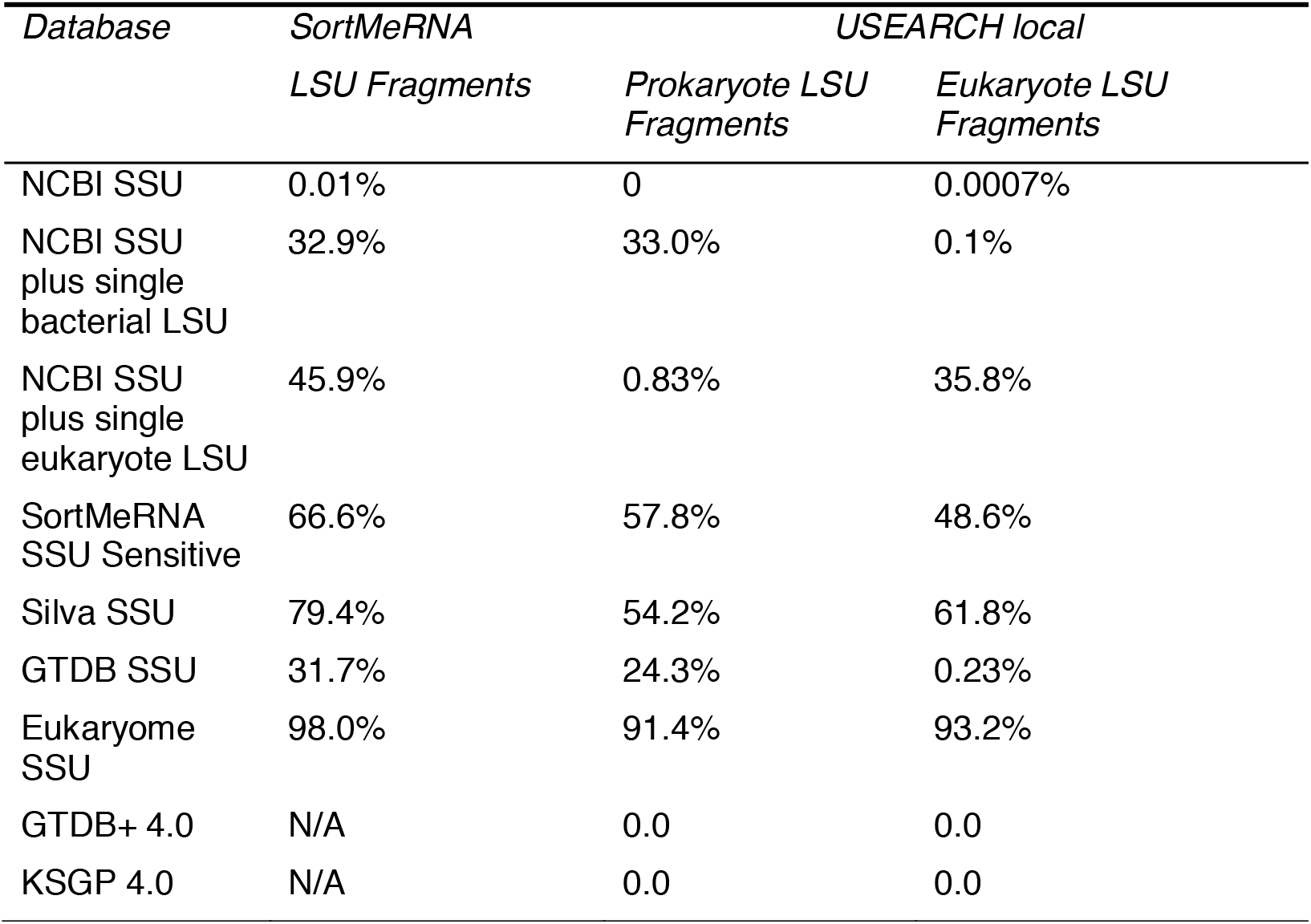
Percentages of 150 bp LSU rRNA fragments erroneously classified as SSU using SortMeRNA and USEARCH local and various databases. SortMeRNA fails when used with large databases such as GTDB+ and KSGP so these results are not available.

### Taxonomic annotation of SSU reads in an example total RNASeq data set

We illustrate these methods using RNASeq data for an estuarine sediment. 8.3 million reads (97% of the total) were retained after quality trimming, approximately half of which represented merged paired end reads. rRNA makes up 80-90% of RNA (12,58), so based on the relative lengths of SSU and LSU sequences, we would expect approximately 35% of total RNA reads to be SSU sequences. In line with this, SortMeRNA using the NCBI SSU database and mTAGs identified 38.5% and 38.3% as SSU respectively (Table 4). USEARCH alignment of all reads to KSGP 4.0 gave a slightly smaller number (38.1%). By contrast, SortMeRNA using its own sensitive database and USEARCH alignment to the SILVA database identified roughly twice as many reads (78.3% and 77.3% respectively) as SSU. However, when LSU contamination was removed from the SILVA and SortMeRNA databases, SortMeRNA and USEARCH identified similar numbers of SSU reads as other approaches (37.9% and 38.6% respectively). mTAGs compares each read with HMMs for SSU and LSU of all three domains of life and chooses the best fitting model, extracting LSU and SSU reads for each domain into separate files. When reads identified as LSU by mTAGs were aligned against SILVA, 3.2 million (9.1%, 68.4% and 69.7% of Archaea, bacteria and eukaryote LSU reads), approximately the same as the number of total reads with matches in SILVA but not in SILVA-cleaned. Only 6 358 LSU reads (0.08% of clean reads) had matches in SILVA-cleaned, falling to 3 428 (0.04% of clean reads) in KSGP 4.0, confirming that the high number of reads identified as SSU when using the raw SILVA and SortMeRNA databases is a consequence of LSU reads being incorrectly matched/extracted in approximately equal numbers to genuine SSU reads. The congruence between alignment and both HMM based analyses when all use rigorously cleaned databases indicates that LSU contamination of annotated sequences can be almost entirely eliminated by using cleaned databases and that for our example data, genuine SSU reads make up approximately 38% of reads. But if LSU contamination is not removed from databases used for SSU extraction and/or alignment, there may be serious consequences for the analysis of real data, potentially leading to LSU sequences making up 50% of the taxonomically assigned supposed SSU reads.

**Table 4.** Numbers of individual reads retained/matched by different methods and databases.

|  | Number | As percentage of clean reads |
| --- | --- | --- |
| Total reads | 8 466 949 |  |
| 100+ bp after cleaning and merging | 8 253 028 |  |
| Identified as SSU by SortMeRNA and NCBI SSU | 3 181 069 | 38.5 |
| Identified as SSU by mTAGs | 3 157 799 | 38.3 |
| Identified as SSU by SortMeRNA with its Sensitive database | 6 465 998 | 78.3 |
| Identified as SSU by SortMeRNA with cleaned Sensitive database | 3 187 519 | 38.6 |
| Identified as SSU by SortMeRNA with combined NCBI and cleaned Sensitive database | 3 188 460 | 38.6 |
| Reads with USEARCH matches to KSGP 4.0 | 3 140 969 | 38.1 |
| Reads with USEARCH matches to raw SILVA 138.1 | 6 376 766 | 77.3 |
| mTAGs LSU Reads with matches to raw SILVA 138.1 | 3 252 093 | 39.4 |
| Reads with USEARCH matches to SILVA-cleaned | 3 130 716 | 37.9 |

We can compare the number of reads assigned to individual phyla using the cleaned and raw SILVA databases (Fig. 1). Read numbers are similar for the majority of phyla, although the doubling of the total number of assigned reads when using contaminated databases means that the apparent relative abundance of most taxa is approximately halved. When the contaminated database is used, the erroneously included LSU reads align with the small number of contaminant sequences and the taxa represented by these can dominate taxonomic assignments. In our example data, this includes reads for which taxonomy was not resolved at all and those identified only as bacteria, which might be mistaken as indicating the presence of common but phylogenetically divergent lineages. The bacterial phyla Elusiomicrobiota and Firmicutes show apparent 255 and 60 fold increases in relative abundance respectively. The proportion of LSU reads matching contaminant sequences could be reduced by using a higher sequence similarity threshold for alignments, but even setting this threshold at 90% similarity would still retain around 20% of LSU reads (Fig. 2). Use of a higher threshold would also mean that some SSU sequences from poorly characterised lineages would be discarded, with greater impacts on Archaea and eukaryotes than on bacteria (see discussion below of Fig. 2). Therefore rigorous cleaning of LSU contamination from databases is preferable.

**Figure 1.**
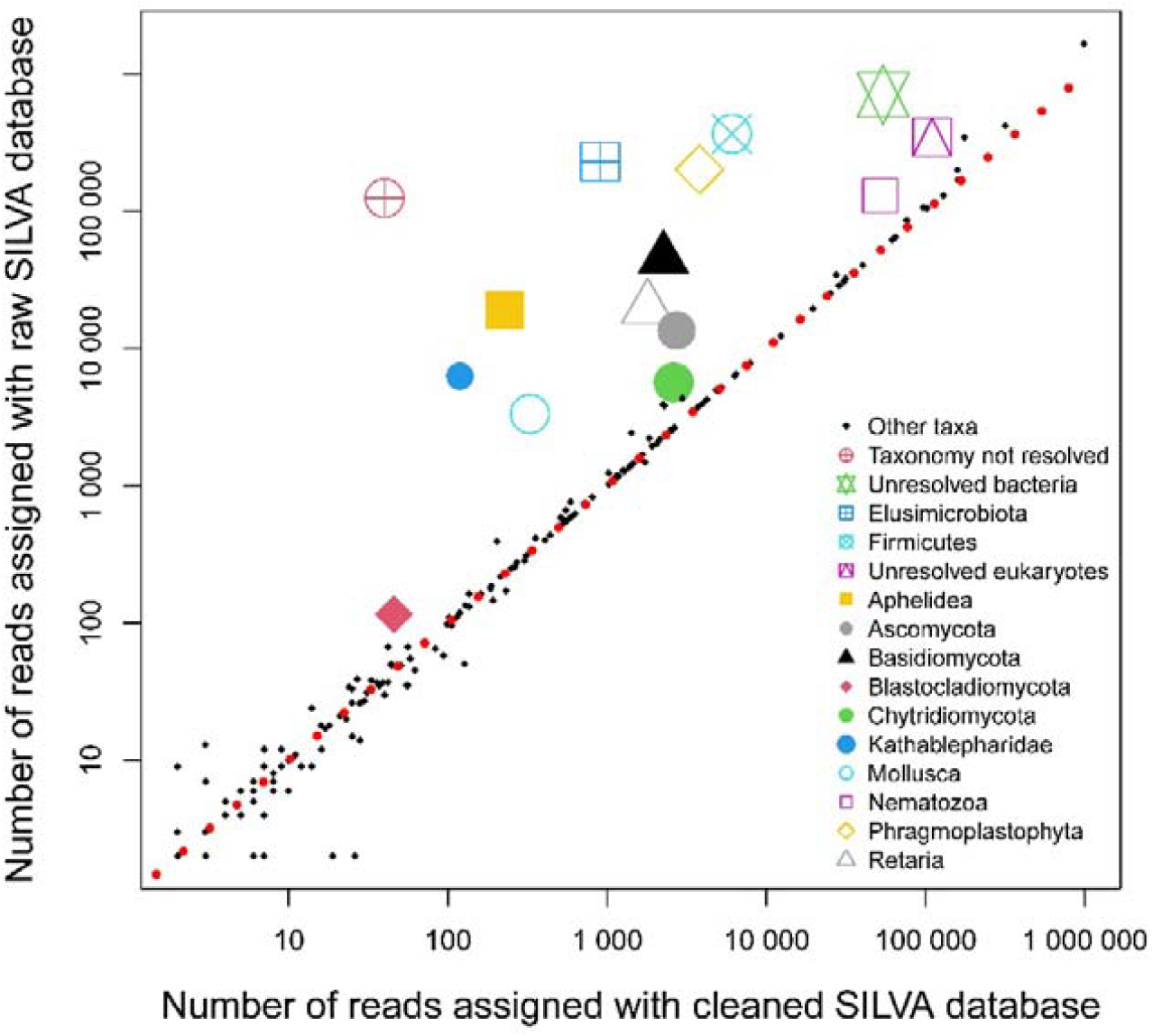
Numbers of reads from example total RNASeq dataset assigned to phylum level taxa using SILVA 138.1 database with (horizontal) and without (vertical) removal of LSU contamination. Highlighted phyla have more than twofold higher counts using the contaminated database. Red dotted line indicates equal read numbers with both versions of SILVA.

**Figure 2.**
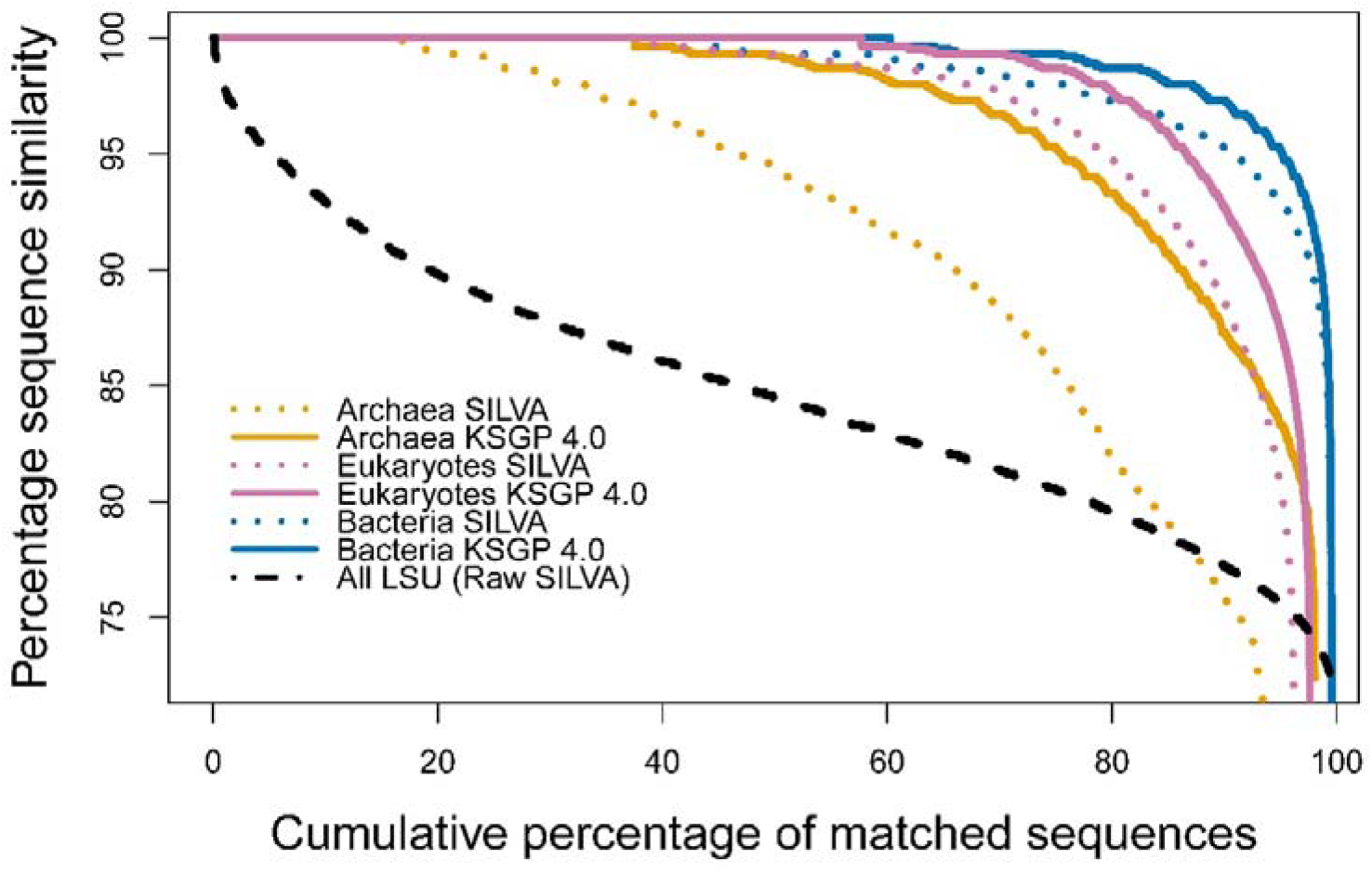
Sequence similarity between individual SSU rRNA reads and best matching database sequence, sorted and plotted in rank order. Dotted lines are the best match in SILVA-cleaned (with LSU contamination removed) and solid lines the best match in KSGP 4.0. Black dashed line shows the cumulative distribution of sequence similarities between individual reads identified as LSU by mTAGs and SILVA 138.1 without removal of LSU contamination.

For more than 99% of reads passing the quality thresholds, SortMeRNA and USEARCH are in agreement on whether or not they should be identified as SSU. A minority of the remainder (0.1% of total reads) have matches in KSGP 4.0 but are not identified as SSU by SortMeRNA. 12% of these aligned with mitochondrial 12S sequences so are appropriately excluded. The great majority of the remainder were identified as eukaryote, including 37% Amoebozoa and 25% Heterolobosa. More than half of the individual pairwise matches for these reads were to sequences in Eukaryome which are derived from collections of long PCR products (identifiers beginning EUK) and a further quarter to sequences from the Karst long environmental RNASeq data. This suggests that there are poorly characterised lineages of unicellular eukaryotes which are absent from the database used by SortMeRNA. The largest group of ambiguous reads (0.5% of total reads) were identified as SSU by SortMeRNA but did not have matches in KSGP 4.0. However, the detailed SortMeRNA BLAST report showed that 91% of these were based on an alignment of less than 100 bp with the SSU database used so would be unlikely to satisfy our minimum alignment length criterion for USEARCH matches even if they were 100% identical to sequences in the database, so are correctly excluded at the alignment stage. The remaining 9% (0.05% of clean reads) may be real SSU sequences that are being overlooked because they have no close matches in KSGP. Approximately half had matches to foraminiferan (Rhizaria) 18S sequences in Genbank and 32% had matches in a collection of long PCR products generated from the same estuarine sediments so this figure will be reduced further as known SSU sequences, particularly from eukaryotic microbes, are added to databases. If initial SSU extraction uses the most comprehensive of our databases, SSU sequences representing approximately 0.08% of clean reads may be overlooked. But these calculations indicate that more than 99.5% of SSU reads will be included in the annotated sequences using our recommended approach. When applied to metagenomic data, approximately 75% of the reads pre-selected by SortMeRNA do not have USEARCH matches in KSGP 4.0 but almost all of these consist of short tandem repeats rather than SSU sequences. This will be explored in detail elsewhere.

We can assess database coverage in more detail by examining the distribution of sequence identities between individual reads and the best database match and can do this separately for each domain of life as mTAGs disaggregates reads by domain (Fig. 2). Comparing the distributions of the closest matches in SILVA-cleaned and KSGP 4.0 shows how much the completeness of taxonomic coverage has been improved by recent database updates. Both KSGP 4.0 and SILVA have matches to 99.5% of sequences identified as bacterial SSU but matches in KSGP 4.0 have somewhat higher identities and the number of exact matches increases to 60% as compared with 42% in SILVA. Slightly more eukaryote reads have matches in KSGP 4.0 than in SILVA (97.6% vs 96.2%), again with more exact matches (58% vs 40%) and somewhat higher identities for the remainder. KSGP 4.0 provides a particularly marked improvement for Archaea with 98.1% having matches (compared to 93.4% in SILVA); 38% exact matches (17% in SILVA) and substantially closer matches for the remainder.

As noted previously (32), the additional sequences derived from Karst *et al. (59)* make substantial improvements in database coverage for Archaea compared to SILVA. Eukaryome’s (31) strategy of extracting sequences of long PCR products is making a substantial contribution to improving database coverage for eukaryotes. But even for bacteria, sequences from Karst *et al*. and GTDB make some improvement in sequence identities between KSGP 4.0 and SILVA.

We have seen in this study that database contamination can have serious impacts on taxonomic assignments. As GreenGenes2 is focused on prokaryote 16S sequences, the overwhelming majority of which are short V4 amplicons (60), the raw SILVA database is widely used in metatranscriptomic and metagenomic based work. As far as we are aware, the phyloFlash pipeline (38) is the only other work to use a database that is rigorously cleaned of LSU contamination. The extent to which database contamination impacts a particular metagenomic or metatranscriptomic study will depend upon the bioinformatic tools used. If SSU pre-selection successfully removes the great majority of LSU reads by, for example, using well designed HMMs such as those within mTAGs, then taxonomic assignments will not be markedly impacted if alignment uses a database contaminated with LSU sequences. However, there are a number of metagenomic studies in the literature that use the SortMeRNA and SILVA databases without removing LSU contamination and will be severely impacted. As this problem is widespread in the literature, we have chosen not to single out particular examples.

### A recommended bioinformatic strategy

SSU pre-selection and sequence alignment can misidentify LSU reads in equal numbers to SSU when used with contaminated databases, so at least one of these, and ideally both, should use databases from which LSU contamination has been removed. A number of alternative programs could be used, but we illustrate our approach using USEARCH and SortMeRNA.

Our preference is to carry out taxonomic assignment by combining local alignment using USEARCH with LCA. Pre-selection of SSU reads using SortMeRNA or another rapid tool can typically be carried out much faster than full sequence alignment. For metagenomic data, SSU sequences may make up only about 0.05% of reads, so for most users, pre-selection of reads before taxonomic assignment is likely to be essential. For total RNASeq data, the proportion of SSU reads is likely to be similar to the 38% in our example dataset, so SSU pre-selection will offer a smaller reduction in overall computation time and might be omitted. However, detailed examination of sequences identified as SSU by one of these approaches but not the other can give insights into the completeness of database taxonomic coverage. Our recommended approach, including pre-selection of SSU reads, is outlined in Box 1. This can be carried out using our total_rnaseq script (see Data Availability), generating a taxonomic annotation for each read and tables of read counts at each taxonomic level.

**Box 1.** Recommended bioinformatic strategy, as implemented in the script total_rnaseq (see data availability).

| Stage of analysis with example tool used | Notes |
| --- | --- |
| 1. Removal of LSU contamination from databases using SortMeRNA and cleaned NCBI LSU database | Already completed for KSGP 4.0; NCBI SSU and SortMeRNA sensitive databases. |
| 2. Read quality screening and adapter removal (Trimmomatic) |  |
| 3. Paired end merging, retaining only R1 if reads do not overlap (USEARCH fastq mergepairs) | Some authors concatenate non-overlapping reads. The variable length gaps in alignments with database sequences will have unpredictable consequences for the LCA algorithm that we use for taxonomic assignment. |
| 4. SSU pre-selection using SortMeRNA | Cleaned NCBI SSU database is small and fast. Cleaned SortMeRNA database is larger and selects a slightly higher number of sequences. The two can be concatenated and further sequences could be added by the user. |
| 5. USEARCH local alignment | Individual SSU reads cover only a small proportion of the full SSU, and not all database sequences are full length, so use of local alignment is essential. This can reduce the effectiveness of k-mer based approaches to speed up database searching (see text). |
| 6. Taxonomic assignment of individual reads using LCA (LCA 0.25 from LotuS2) | KSGP 4.0 comes with three different taxonomy files. We recommend KSGP + For the exploration of poorly characterised taxonomic lineages while KSGP LCA provides the most conservative taxonomic annotations. |
| 7. Constructing tables of taxon counts at each taxonomic level. |  |

The user can alter the databases used for pre-selection and alignment. Pre-selection using a small but clean SSU database is computationally efficient and minimises the extent of LSU contamination. However, this carries with it a risk of incorrectly discarding SSU reads from poorly known taxonomic groups that do not have a match in the database. When SortMeRNA is used with the cleaned SortMeRNA sensitive database, it identifies 0.1% more reads as SSU than when used with the substantially smaller NCBI SSU database. If an alternative tool is used for pre-selection, the user should check that it correctly selects SSU fragments and rejects LSU fragments using the files we have created from the NCBI SSU and LSU databases.

For a subset of samples, it is valuable to compare ambiguities between the results of SSU pre-selection with alignment of all reads. This will identify whether there are any taxonomic lineages which are missed in the pre-selection or are absent from the database used for taxonomic alignment. SortMeRNA constructs its HMM at run-time from a database of sequences, which makes it straightforward to screen the SSU database for LSU contamination and to improve taxonomic coverage of pre-selection by adding in additional sequences from either KSGP 4.0 or elsewhere. Most other HMM based tools provide pre-constructed HMMs. In some cases, the sequence alignments on which these are based are available, but this is often not the case, making removing contaminants or adding sequences to increase taxonomic coverage difficult or impossible. If taxonomic gaps in databases become apparent, sequences from NCBI nt, long PCR products amplified from the study sites or other sources could be added in to the pre-selection and/or the taxonomic database. If they are being added to the taxonomic database, they will need to be annotated manually or using the approaches on which KSGP is based.

We have not included the mTAGs package in this pipeline, even though it has some valuable features. The initial separation of SSU sequences within mTAGs is based on finding the best fitting HMM from six that are constructed separately for both SSU and LSU of Bacteria, Eukarya and Archaea. This leads to lower rate of false positives than using SortMeRNA with a single SSU database and allows separate assessment for the three domains of completeness of the taxonomic database. However, it fails to detect a number of SSU reads, presumably reflecting incomplete taxonomic coverage by its databases. As information on its HMMs is not available, it is not possible to modify this. In addition, mTAGs own taxonomic assignment procedure is much less effective than our LCA based approach and it replaces sequence read identifiers with its own labels, which makes some downstream analyses more challenging.

## Acknowledgements

The bioinformatic analysis was carried out on the High Performance Computing Cluster supported by the Research and Specialist Computing Support service at the University of East Anglia. We are grateful to Robert Edgar for helpful discussions on USEARCH and other alternatives to BLASTN.

## Author contributions

AG designed the study, analysed the data and wrote the first draft of the manuscript. CSD carried out the laboratory work and revised the manuscript.

## Data availability

KSGP and GTDB+ 4.0 are available at https://ksgp.earlham.ac.uk/. A script to carry out the analysis in Box 1, total_rnaseq and the databases required to run this are available at https://github.com/AGrantUEA/total_RNASeq. The repository also contains the other databases used in this work (see table 1) and the scripts used in this paper. The example RNASeq data are deposited in ENA under project PRJEB116069.

